# A Dual Effort Choice paradigm reveals pharmacological divergence between intrinsic and extrinsic motivation within the same task environment

**DOI:** 10.64898/2026.11.12.766115

**Authors:** Froso Fachouridou, Foteini Xeni, Caterina Marangoni, Peter Salisbury, Eleanor W Grayson, Emma S J Robinson, Megan G Jackson

**Affiliations:** School of Psychology and Neuroscience, Biomedical Sciences Building, University of Bristol, Bristol, England

## Abstract

Motivation loss is a transdiagnostic symptom, prevalent across neuropsychiatric and neurodegenerative diseases. Current treatments fail to effectively treat this symptom domain, giving it significant clinical importance. The fields of neuroscience and drug development have primarily focused on extrinsic motivation, whereby an animal must perform an action to receive an external food or water-based reward. However, many of our everyday activities are facilitated by intrinsic motivation, where completing an action is inherently rewarding. By utilising tasks focused only on extrinsic motivation, we may miss critical neurobiological and therapeutic differences between motivation types. Previous work has shown divergence in drug effect between extrinsic and intrinsic motivation tasks across a wide range of clinically-used drugs, indicating translationally-relevant differences across motivation domains that warrant further study. Therefore, the aim of this work was first to develop a task capable of simultaneously quantifying intrinsic and extrinsically motivated behaviour within the same task environment. Secondly, to use this task to further investigate this neurochemical divergence by examining a wide range of clinically-relevant drugs within this new task context. Here, we find that a pharmacological reduction in dopaminergic signalling reduced motivation across both extrinsic and intrinsic options within the new task, though this effect was mechanism dependent. In contrast, acute serotonergic and dopaminergic signalling increase produced a more heterogenous pattern, indicating nuanced mechanistic differences even when presented in the same task environment. Therefore, this task provides a new approach for characterising drug effects on distinct forms of motivated behaviour that may go unnoticed using conventional assays.

## Introduction

Motivation loss is a transdiagnostic symptom, prevalent across a range of neuropsychiatric and neurodegenerative disorders, including Major Depressive Disorder (MDD), Dementia and Schizophrenia^1^. First-line treatments for these disorders including selective serotonin reuptake inhibitors^2,3^, dopaminergic antipsychotics^4–6^ and cognitive enhancers^7^ fail to effectively improve motivation loss and critically, can worsen it. Motivation loss therefore remains very difficult to treat and there have been no advances in terms of novel pharmacological therapies, potentially due to limitations of current animal models

Fundamental neuroscience and drug development have mostly centred around extrinsic motivation, whereby an animal must perform an effortful instrumental response such as nose poking to receive an external reward, such as food or water. Effort Based Decision Making (EBDM) paradigms such as the Effort for Reward (EfR) task^8^ and Progressive Ratio task (PR)^9^ have advanced our understanding of the neurobiology of extrinsic motivation and have been used to screen a wide range of compounds for therapeutic effect^10–12^. However, humans and non-human animals engage in behaviours that do not require tangible external reward. Many of our everyday activities, such as engaging in hobbies are intrinsically motivated, where engaging in the action itself is inherently rewarding^13^. Non-human animals also engage in intrinsically motivated behaviours, including novelty exploration, puzzle solving and some foraging behaviours^14–16^.

A neurofunctional distinction between extrinsic and natural rewards has been reported in patients with MDD^17^ and although results are mixed, other neuroimaging studies found distinct patterns of regional activation between intrinsic and extrinsically motivated tasks, particularly within frontal and striatal regions^13^. This work provides initial evidence that intrinsic motivation is partially neurobiologically distinct from extrinsic motivation. Assessing this distinction in animal models could therefore improve our understanding of its underlying mechanisms and identify novel therapeutic targets. However preclinical tasks rarely retain or explicitly assess intrinsic motivation. While many tasks are rooted in intrinsic motivation (the initial drive to explore the task environment), it is often rapidly shifted to extrinsic motivation through food restriction and weeks-months of conditioning training. While this facilitates high levels of control over the behaviour, in choosing to focus on only one dimension of motivation, we may miss differences in therapeutic sensitivity and efficacy between motivation types that have significant translational implications for treatment.

The Effort Based Forage (EBF) task^14^ draws on the mouse’s natural drive to forage for nesting material. Foraging output is sensitive to effort requirement but persists even when the mouse is sufficiently warm or has free access to nesting material. Therefore, this task was found to be a uniquely useful measure of intrinsic motivational state. In recent work, it was found that the EBF task showed a profound and consistent divergence in serotonergic antidepressant effect when compared to a typical extrinsically motivated EfR task. In this paradigm, the mouse can choose to complete four nose pokes to receive a palatable reward (a high effort, high reward option), or consume freely consumable chow (a low effort, low reward option). While acute administration of these compounds dose-dependently increased motivation in the EfR task, it decreased motivation in the EBF task^18^. Use of dopaminergic compounds haloperidol and methylphenidate showed a more convergent profile across tasks, while amphetamine produced a similar serotonergic-like divergence^14,19^. These results suggest that both forms of motivation engage partially distinct neurobiological mechanisms with some convergence in the mesolimbic dopaminergic system. The ability to simultaneously investigate the effects of different neurochemical manipulations on both types of motivation within the same test session could enable a more detailed understanding of these differences. However, a task that facilitates the simultaneous measurement of both types of motivation does not yet exist.

Therefore, this work had two core aims. The first was to develop a task capable of simultaneously quantifying intrinsic and extrinsically motivated behaviour within the same task environment, drawing on key elements of the EBF and EfR tasks. The second was to use this task to further investigate the neurochemical divergence in the EBF and EfR tasks by examining the effects of a wide range of clinically relevant drugs within this new task context. We hypothesised that previously observed divergence in drug effect would persist in this novel task, providing further evidence that intrinsic and extrinsic motivation are meaningfully distinct and that both should be included in the pipeline when investigating drug effect. We further hypothesised that individual drugs would produce distinct patterns of effects across the two behaviours, which could provide initial insight into the neurobiological mechanisms underlying their regulation.

## Methods

Four cohorts of C57bl/6j mice (half male, half female, Envigo, UK) were used in these experiments (see **S1** for summary). Three cohorts of mice were housed in enriched (**S2**) open-top Techniplast 1284 conventional caging and one cohort were housed in larger enriched Techniplast 1290 conventional caging (**S2**). Males were housed alone to minimise psychosocial stress associated with aggression and to maintain a positive affective state^20^. Female mice were housed in pairs. Mice underwent twice daily health checks to maintain good welfare. Mice were kept in temperature-controlled rooms (∼21°C) and on a 12:12 reversed light cycle, where lights turned OFF at 8.15am and ON at 20.15pm. All experiments took place in the dark phase during the animals’ active period. Mice were provided food (Purina, UK) and water *ad libitum* including during testing. Before onset of experimentation, mice were habituated to cup handling in accordance with 3Hs principles^21^. Except for experiment 1 which was exploratory, expected effect sizes were drawn from previous work using similar paradigms and pharmacological manipulations, with consideration for potential attrition. In all drug studies, the experimenter was blind to dose, and mice received doses in a randomised, counterbalanced order to mitigate order-of-testing effects. All experiments were conducted in accordance with the Animals Scientific Procedures Act (United Kingdom, 1986) and were approved by the University of Bristol Animal Welfare and Ethical Review Body (AWERB).

### The Dual Effort Choice task

#### Arena

An arena was designed in-house by MGJ to recapitulate principles of the Effort Based Forage task^22,23^ and Effort for Reward task^24^ within the same task environment. The apparatus comprises a safe and enclosed home-like environment which provides shelter and free access to food and water. Two tubes extend from the home area to two separate environments. One contains a nesting material foraging option, where mice can exert effort to obtain Sizzlenest (Datesand, UK) via apertures in a custom designed nesting box (**S3**). Custom components are available for print here https://github.com/meganjackson13/Bedding-box-3D-files. The other environment contains a Feeding Experimental Device 3 (FED3)^25^, a portable open-access pellet dispenser consisting of two nose poke apertures either side of a centrally located magazine. Here, the mice can learn to nose poke to receive a palatable food reward (20mg reward pellet, TestDiet)(**S4**).

#### Habituation

On the first session, mice were put individually into empty arenas containing woodchip for 15 minutes, then 10 minutes on the second and third sessions. The fourth session comprised four hours with access to food and water within the home area. In one of the compartments, a nesting material box was fixed to the back wall where nesting material (Sizzlenest) could be accessed through the widest (easiest) face plate apertures (1.5cm^2^). In the other compartment, a FED device was placed at the back and protected by a Perspex screen, leaving the nose poke apertures and food magazine exposed. Option position was counterbalanced across mice. The FED device was set to Fixed Ratio (FR1), where one nose poke in the active aperture resulted in the delivery of a reward. Position of the active nose poke was also counterbalanced across mice.

Nesting material box weight was measured pre- and post-session, providing the total nesting material foraged output measure. The amount of nesting material dropped in the compartment was also measured. From this, the nesting material shuttled measure was determined (total nesting material foraged – total left in compartment), which represents how much the mouse foraged and carried through to the home area. Amount of nesting material shuttled was expressed as a percentage proportion of total foraged (nesting material shuttled/total nesting material shuttled)*100. The number of reward pellets obtained at the FED device was also recorded.

### Experiment 1: the effect of effort and reward value modulation on choice motivation

#### Training

Mice next completed a 2hr session using the moderately difficult apertures (1.0cm^2^). 2hr sessions were repeated until mice were performing stably over 30 trials for at least three sessions. Training in foraging was not required as it is an innately expressed behaviour, however cohort stability was monitored.

#### Reward devaluation via pre-feeding

10 mins preceding onset of the task, half of the mice were provided with free access to reward pellets within a custom-made bowl and half of the mice were presented with an empty bowl. The bowl was weighed to confirm that mice had consumed pellets. Mice then underwent the DEC task for 2hrs across two sessions in a counterbalanced design.

#### Reward value potentiation via food restriction

2hrs preceding the task, half of the mice had their home cage food removed across two sessions in a counterbalanced design.

#### Foraging effort modulation via aperture change

In the first session, half of the mice had the difficult aperture in the nesting material box (0.75cm^2^), while half had the moderate aperture (1.0cm^2^). This was counterbalanced across two sessions.

#### FED effort modulation via FR increase

Mice first underwent an incremental baseline session of FR3 before increasing to FR5. Mice underwent 4 sessions for FR5 testing before study onset. In a test session, half the mice underwent FR5 testing and half at FR1, counterbalanced across two test sessions. Each test session was preceded by a FR5 baseline session.

### Experiment 2: acute pharmacology in the DEC task

#### Training

Following habituation, mice completed a 2hr session with the moderately difficult apertures (1.0cm^2^). Mice ascended through FR1, FR2 and FR4 if they completed a session with 30+ trials. Stability in performance across both measures was monitored before onset of testing.

Three cohorts of 16 mice were used for the pharmacology studies. Doses/pre-treatment times were selected based on the minimally effective dose that showed a divergent effect in the Effort Based Forage task and Effort for Reward task in previous work. Where minimally effective doses differed between tasks, a dose response curve was conducted instead. In previous work, dose ranges were based on human dose ranges using the animal-equivalent dose equation and aligned to clinically relevant doses. Pretreatment times (t-) were based on T_max_^14,18^.

Pre-treatment times, vehicles, route of administration and doses are outlined in the table below (**table 1**). Drugs were administered at a dose volume of 10ml/kg. Mice were trained to voluntarily consume the drug in a palatable solution to reduce stress associated with dosing and maximise translation to human methods of consumption (**S5**). Drugs were administered sub-cutaneously if the drugs had low oral bioavailability. In this instance, mice were trained in a lower restraint method for dosing^21^ (**S6**).

| Drug | Pre-treatment time | Dose (mg/kg) | Route of administration | Vehicle | Supplier |
|---|---|---|---|---|---|
| Escitalopram | 1hr | 3 | Oral | Water + condensed milk (20%) | Lundbeck ltd |
| Sertraline | 4hr | 3, 10 | Oral | 1% DMSO, 2% cremophore, 97% water, condensed milk (50%) | HelloBio |
| Venlafaxine | 1hr | 10 | Oral | Water + condensed milk (20%) | Rosemont |
| Methylphenidate | 15 min | 1, 3,10 | Oral | Water + condensed milk (20%) | Sigma UK |
| Amphetamine | 30 min | 0.3 | Sub-cut | Saline | Sigma UK |
| Tetrabenazine | 2hrs | 6 | Sub-cut | 1% DMSO, 2% cremophore, 97% saline |  |
| Haloperidol | 1hr | 0.1 | Sub-cut | 1% DMSO, 2% cremophore, 97% saline | Sigma UK |
| Pramipexole | 1hr | 0.1 | Oral | Water + condensed milk (20%) | HelloBio |

#### Testing

A test session was 2hrs long, consisting of the moderate aperture on the foraging option, and an FR4 requirement for the FED option. Test sessions were at least two days apart to allow sufficient drug wash out between sessions. Mice were given the drugs in a within-subject, counterbalanced design so that all mice received each dose.

### Statistics

Data were tested for normality using the Shapiro-Wilk test. Where data were normally distributed, a paired t-test was used for vehicle vs single dose studies or a repeated measures ANOVA for multi-dose experiments. If data were non-normally distributed, a Mann-Whitney U or Friedman’s test was used instead. The threshold for significance was set at p<0.05. Outliers were defined as values 2±SDs away from the mean. In paired comparisons, outliers were excluded. In repeated measures comparisons, outliers were replaced with the group mean to permit repeated-measure analysis. A table summarising all exclusions is summarised in **S7**. Data were analysed and graphed using Graphpad Prism v10.0.3.

## Results

### Foraging and instrumental responding can be independently modulated within the DEC task environment

Food restriction had no effect on foraging behaviour (Fig.**1A-C**) but did increase the number of trials completed (t_(5)_=5.665, p=0.0024) (**Fig.1D**). Pre-feeding had no effect on the foraging behaviour (**Fig.1E-G**) but decreased number of trials completed (t_(5)_=3.967, p=0.0107) (**Fig.1.H**).

**Fig. 1.**
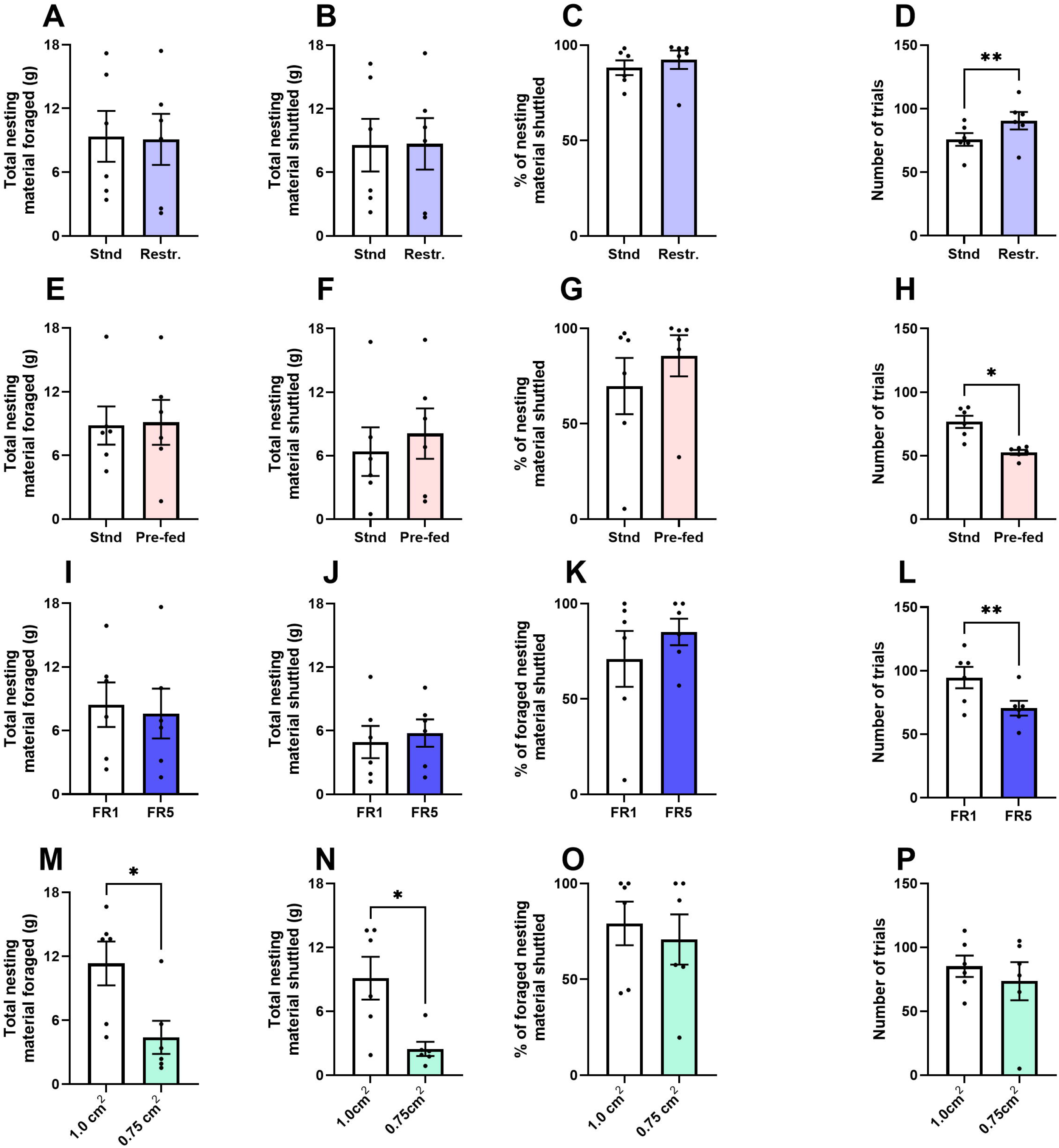
Foraging and instrumental responding do not influence each other within the DEC task environment. (**A-C**) Food restriction 2-hours prior to access to the task did not influence foraging behaviour. (**D**) However, number of trials increased (p<0.01). (**E-G**) Pre-feeding mice with pellets prior to running the task did not influence foraging behaviour. (**H**) However, de-valuing pellets decreased number of trials (p<0.05). (**I-K**) Increasing the effort of conditioned reinforcement behaviour from FR 1 to 5 did not alter foraging behaviour, and (**L**) resulted in lower trials (p<0.01). (**M-N**) Increasing the effort required to pull nesting material out of the box decreased total nesting material foraged and shuttled (p<0.05), (**O**) with no effect on % nesting material foraged, or (**P**) the number of trials completed. Error bars are mean ±SEM, *p<0.05, **p<0.01.

Increasing the effort of instrumental responding from FR1 to 5 did not alter foraging behaviour (**Fig.1I&K**) but did decrease the number of trials completed (t_(5)_=5.245, p=0.0033) (**Fig.1L**). Increasing the effort required to forage decreased total nesting material foraged (t_(5)_=3.945, p=0.0109) (**Fig.1M**) and total nesting material shuttled (t_(5)_=3.805, p=0.0126) (**Fig.1N**). There was no effect of decreasing the aperture size on % nesting material shuttled or number of trials completed (**Fig.1O&P**).

### Acute antidepressant treatment exerts distinct effects on foraging and instrumental responding in the DEC task

3mg/kg escitalopram reduced total nesting material foraged (t_(13)_=2.212, p=0.0456) (**Fig.2A**) but not nesting material shuttled or % nesting material shuttled (**Fig.2B&C**). It increased the number of trials completed relative to the vehicle (t_(12)_=3.982, p=0.0018) (**Fig.2D**).

**Fig. 2.**
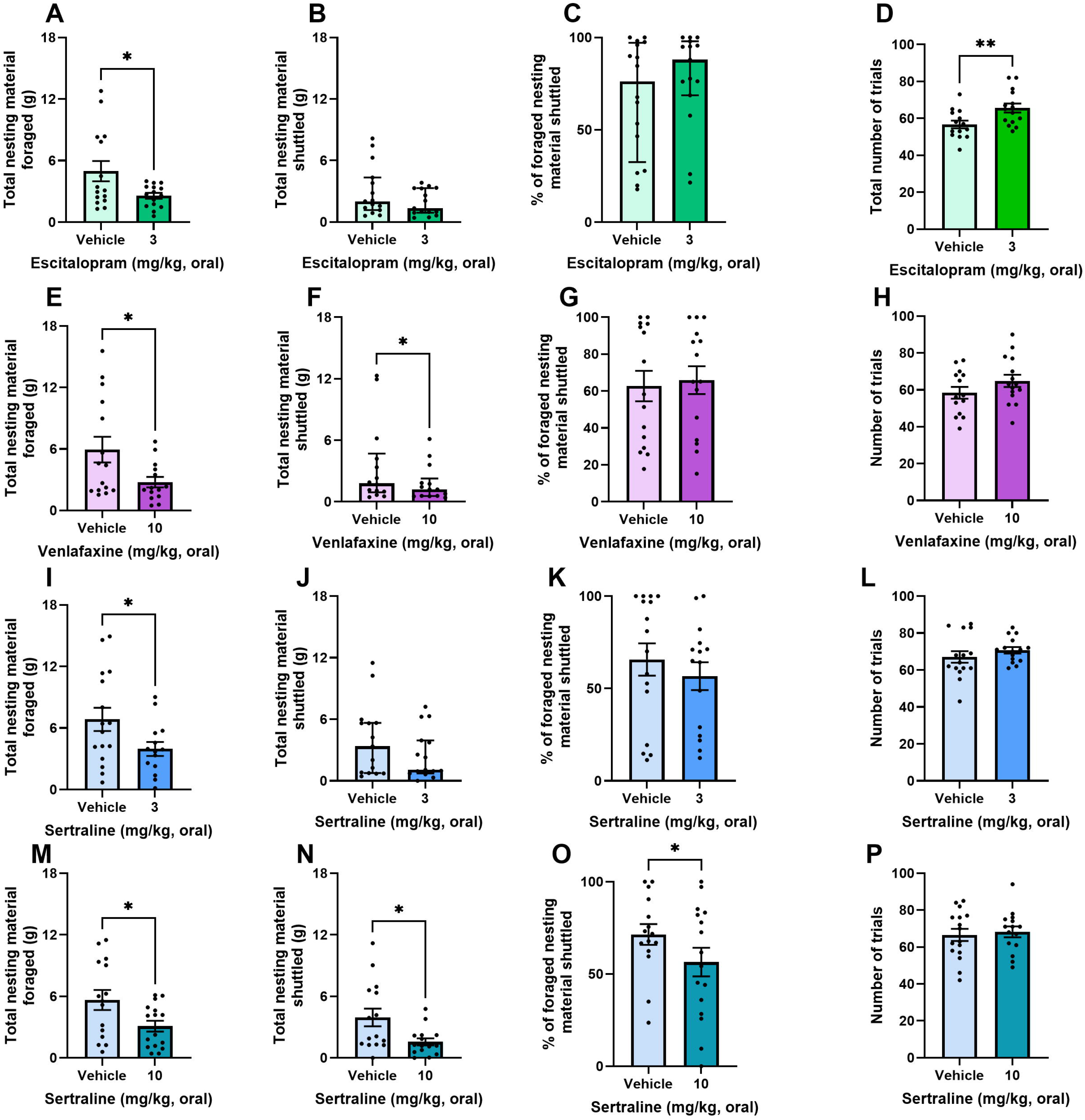
Acute antidepressant treatment exerts distinct effects on foraging and instrumental responding in the DEC task. Mice were dosed with 3mg/kg escitalopram and underwent the DEC task (**A-D**). (**A**) 3mg/kg escitalopram reduced total nesting material foraged (p<0.05), without effect on nesting material shuttled or %nesting material shuttled (**B, C**). (**D**) 3mg/kg escitalopram increased the number of trials completed (p<0.01). Mice were dosed with 10mg/kg venlafaxine and underwent the DEC task (**E-H**). (**E**) 10mg/kg venlafaxine reduced total nesting material foraged (p<0.05), (**F**) and nesting material shuttled (p<0.05). Venlafaxine had no effect on % nesting material shuttled, or number of trials completed (**G, H**). Mice were dosed with 3mg/kg sertraline and underwent the DEC task (**I-L**). (**I**) 3mg/kg sertraline reduced total nesting material foraged (p<0.05). 3mg/kg sertraline had no effect on total nesting material shuttled (**J**), % nesting material shuttled (**K**), or number of trials completed (**L**). Mice were dosed with 10mg/kg sertraline and underwent the DEC task (**M-P**). (**I**) 10mg/kg sertraline reduced total nesting material foraged (p<0.05), total nesting material shuttled (p<0.05) (**N**), and % nesting material shuttled (p<0.05) (**O**). There was no effect on number of trials completed (**P**). Bars are mean ± SEM or median ± interquartile range where data are non-parametric, *p<0.05, **p<0.01.

10mg/kg venlafaxine reduced nesting material foraged (t_(13)_=2.612, p=0.0215) (**Fig.2E**) and nesting material shuttled (W=-57.00, p=0.0479) (**Fig.2F**). There was no effect of venlafaxine on % nesting material shuttled, or number of trials completed (**Fig. 2G&H**).

Both 3 and 10mg/kg sertraline reduced total nesting material foraged (t_(13)_=2.791, p=0.0153, t_(14)_=2.888, p=0.0119 respectively) (**Fig.2I&M**). 10mg/kg sertraline reduced nesting material shuttled (t_(14)_=2.957, p=0.0104), with a trend at 3mg/kg (W=-55, p=0.0906) (**Fig.2N&J**). 3mg/kg had no effect on % shuttled (**Fig.2K**), while 10mg/kg sertraline caused a reduction on % nesting material shuttled (t_(14)_=2.337, p=0.0348) (**Fig.2O**). Neither dose affected trials completed (**Fig.2L&P**).

### Foraging behaviour and instrumental responding are differently affected by dopamine potentiators

0.3mg/kg amphetamine had no effect on total nesting material foraged or nesting material shuttled (**Fig.3A&B**), however there was a trend towards decreased % nesting material shuttled (t_(15)_=1.896, p=0.074) (**Fig.3C**). There was no effect of 0.3mg/kg amphetamine on number of trials completed (**Fig.3D**). To assess a potential interaction between drug effect and intrinsic motivational state, the cohort was then stratified into ‘high’ and ‘low’ performers using a median split of nesting material shuttled under vehicle conditions. There was a drug*motivational state interaction (F_(1,14)_=26.62, p=0.0001) on nesting material foraged where amphetamine decreased nesting material foraged in the higher motivational group (p=0.0005), with a trend towards increase in the lower motivational group (p=0.0555) (**Fig.3E**). There was also a drug*motivational state interaction effect (F_(1,13)_=7.326, p=0.0180) on nesting material shuttled. Amphetamine decreased nesting material shuttled in the higher motivational group (p=0.0171), with no effect in the lower motivational group (**Fig.3F**). There was no drug*motivational state interaction effect on trial number (**Fig.3G**).

**Fig. 3.**
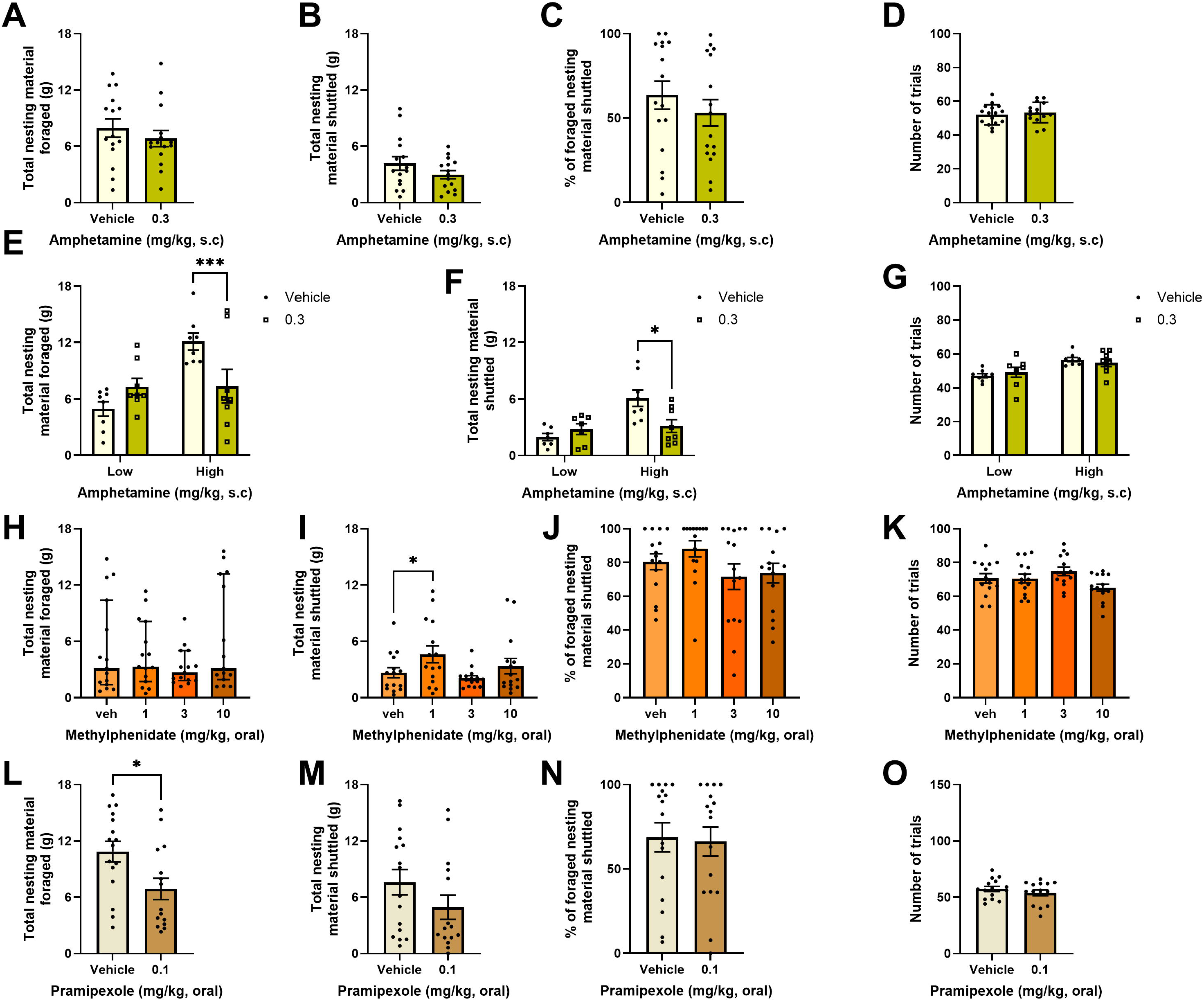
Foraging behaviour and instrumental responding are differently affected by dopamine potentiators. Mice were dosed with 0.3mg/kg amphetamine and underwent the DEC task (**A-D**). 0.3mg/kg amphetamine had no effect on foraging behaviour (**A-C**) or number of trials completed (**D**). 0.3mg/kg amphetamine reduced total nesting material foraged, and nesting material shuttled on high performers only (**E, F**). There was no effect of the drug on number of trials completed in either group (**G**). Mice were dosed with 1-10mg/kg methylphenidate and underwent the DEC task (**H-K**). There was no effect of the drug on total nesting material foraged (**H**). 1mg/kg methylphenidate increased total nesting material shuttled (p<0.05) (**I**), with no effect on % nesting material shuttled (**J**) and number of trials completed (**K**). Bars are mean ± SEM or median ± interquartile range where data are non-parametric *p<0.05, ***p<0.001.

There was no effect of methylphenidate on total nesting material foraged (**Fig.3H**) but there was on nesting material shuttled (F_(2.252,31.53)_=3.934, p=0.0257), where mice treated with 1mg/kg methylphenidate shuttled more nesting material back to the main box compared to the vehicle (p=0.0168) (**Fig.3I**). There was no effect of the drug on % nesting material shuttled (**Fig.3J)**. There was a main effect of the drug on trial number (F_(2784, 37.12)_ =3.221, p=0.0366), however this did not reach post-hoc significance (**Fig.3K**).

A dose response for pramipexole (0.001-0.1mg/kg) was generated in the EBF task (**S8**), allowing selection of an appropriate dose for the DEC task (0.1mg/kg). 0.1mg/kg pramipexole reduced nesting material foraged (t_(14)_=2.501, p=0.0254) (**Fig.4L**). There was no effect of the drug on nesting material shuttled and % nesting material shuttled (**Fig.4M&N**). There was a trend towards a decrease in trials number (t_(13)_=1.870, p=0.0841) (**Fig.4O**).

**Fig. 4.**
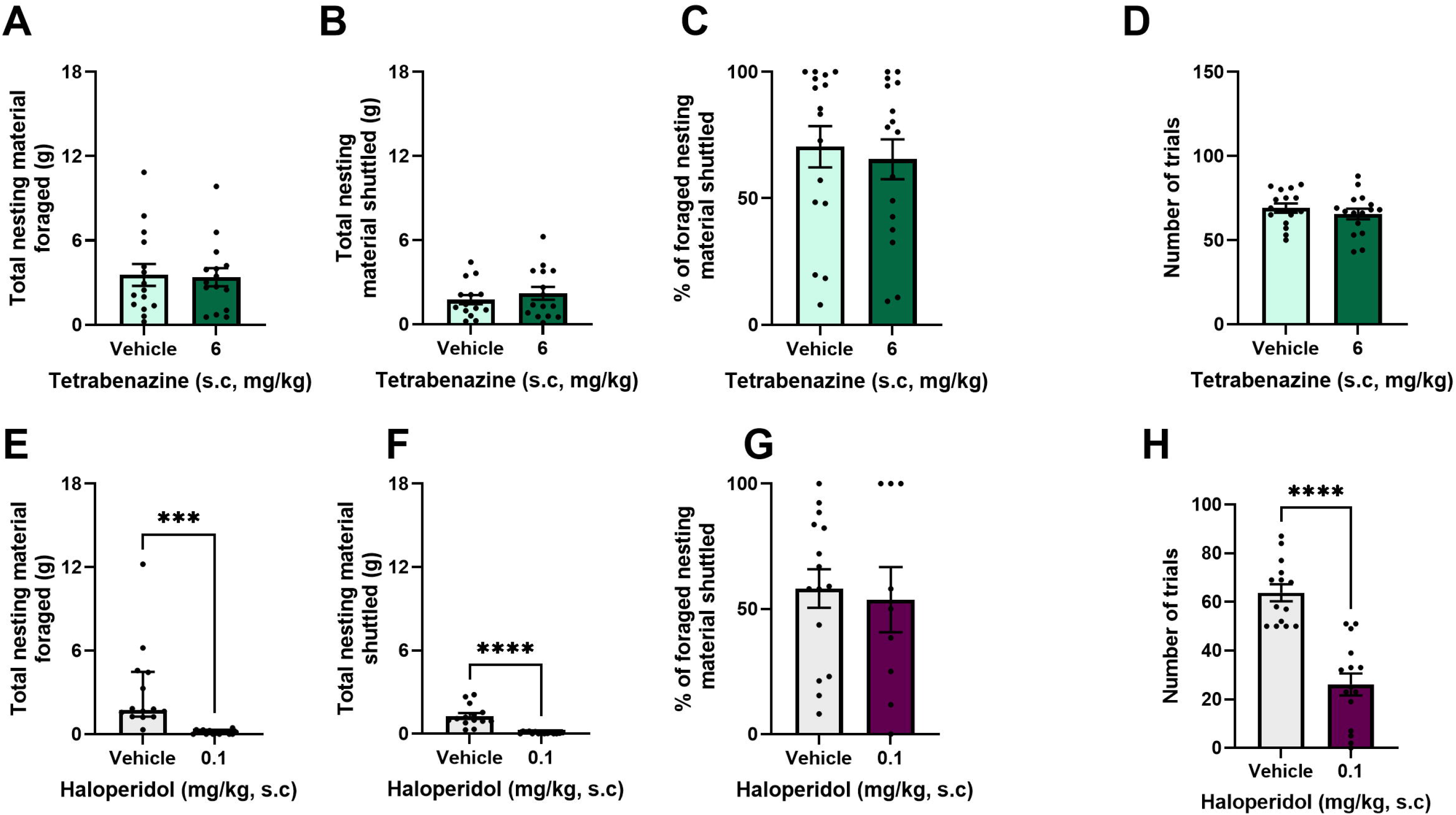
Dopamine inhibition via haloperidol but not tetrabenazine supresses task performance. Mice were dosed with 6mg/kg tetrabenazine and underwent the DEC task (**A-D**). 6mg/kg tetrabenazine had no effect on foraging behaviour (**A-C**) or number of trials completed (**D**). Mice were dosed with 0.1mg/kg haloperidol and underwent the DEC task (**E-H**). 0.1mg/kg haloperidol reduced total nesting material foraged (p<0.001) (**E**), and nesting material shuttled to the main box (p<0.0001) (**F**), with no effect on % nesting material shuttled (**G**). 0.1mg/kg haloperidol reduced number of trials completed (p<0.0001) (**H**). Bars are mean ± SEM or median ± interquartile range where data are non-parametric *p<0.05, ***p<0.001, ****p<0.0001.

### Dopamine inhibition via haloperidol but not tetrabenazine supresses task performance

A dose response for tetrabenazine (2-8mg/kg) was generated in the EBF task (**S9**), allowing selection of an appropriate dose for the DEC task that impaired foraging but did not change general movement in the foraging area (6mg/kg). 6mg/kg tetrabenazine did not affect foraging behaviour or number of trials completed (**Fig.4A-D**). 0.1mg/kg haloperidol reduced total nesting material foraged (W=-105.00, p=0.0001) (**Fig.4E**) and total nesting material shuttled to the main box (t_(13)_=5.846, p>0.0001) (**Fig.4F**). There was no effect of the drug on % nesting material shuttled (**Fig.4G**). 0.1mg/kg haloperidol reduced number of trials completed (t_(13)_=7.234, p<0.0001) (**Fig.4H**).

## Discussion

Bidirectional alteration of reward pellet value via food restriction and pre-feeding altered number of trials but did not affect foraging output. Similarly, increasing effort by increasing the FR requirement reduced number of trials but did not affect foraging. Increasing foraging effort by reducing aperture size reduced foraging but did not affect number of trials. These findings suggest that each behaviour can be independently modulated within the DEC task environment. We next found that the antidepressants escitalopram, venlafaxine and sertraline induced a deficit in foraging consistent with previous work but only escitalopram increased number of trials. Dopamine agonists produced varied responses in foraging but not in trials completed. The stimulant amphetamine impaired foraging but only in a highly motivated sub-population of mice. The dopamine reuptake inhibitor methylphenidate dose-dependently increased foraging while the D_2_/D_3_ receptor agonist pramipexole reduced foraging. Dopamine antagonists had a more similar effect across behavioural options. The VMAT2 inhibitor tetrabenazine had no effect on either foraging or number of trials, while D_2_ antagonist haloperidol impaired both foraging and number of trials. The following discussion will consider how this pattern of results advances our understanding of intrinsic and extrinsic motivation and the utility of the novel DEC task in assessing these differences.

### The DEC task environment facilitates independent modulation of intrinsic and extrinsic motivation

Goal-directed behaviour driven by extrinsic reinforcement such as food delivery is sensitive to its subjective value^26^, and the effort required to obtain it^27^. Increasing reward value via food restriction increased responding for food reward at the FED device while pre-feeding^28,29^ reduced responding. Notably, modulation of food reward value did not impact foraging, indicating that the motivational value of these behavioural options is distinct. Increasing effort via the FR requirement reduced trial number as previously reported^30,31^ and did not affect foraging. Together these findings demonstrate that extrinsic goal-directed behaviour is successfully measured in this arm of the DEC task and in a manner distinct from the foraging option.

As intrinsically motivated behaviours are not driven by external reinforcement, effort requirement for foraging was increased. Foraging was reduced as previously observed^14^, while number of trials were unaffected. % nesting material shuttled, regarded as a measure of anxiety-relevant behaviour^14^ remained unaltered across manipulations. Together, these findings establish that the two behavioural options within the DEC task are responsive to changes in their respective costs and values and demonstrates that suppression of one behaviour does not necessarily enhance the other.

### Pharmacological heterogeneity across foraging and instrumental responding options

Assessing serotonergic function in the context of motivation has significant clinical importance due to the limited efficacy of serotonergic antidepressants for this symptom domain^2^. Utilising the minimally effective doses that induced change in motivational state independent of motoric changes^18^, we found that a divergence in antidepressant effect persists in the DEC task environment, largely driven by a deficit in total nesting material foraged versus a relative lack of effect on number of trials at the FED device. Notably, lack of effect on trials contrasts with findings from our previous work with the EfR task^18^. Effects may be more pronounced in the conventional EfR task where behaviour is guided through external control of motivation via food restriction and a simplified testing environment. This suggests that under conditions of free feeding and environmental complexity, extrinsic motivation is less affected by acute serotonergic increase. This demonstrates that drug effect is dependent not just on the behavioural action itself but the environment in which it is set. However, foraging remains vulnerable, indicating this effect persists even under greater task complexity and is not linked to general increase in anxiety behaviour.

Contrasting with the other compounds, escitalopram increased trial number in addition to supressing foraging, showing a more marked divergence. It is currently unclear what drives this difference but may relate to its greater selectivity for SERT^32^. To test whether lack of effect of sertraline on trials was driven by a comparatively lower SERT occupancy at the chosen dose, a higher dose was tested. Lack of effect at the higher dose suggests the differences are more likely driven by variances in compound binding profiles. Further investigation is required to understand the nature of these differences but may have important clinical implications under conditions of chronic dosing. While total foraging levels were suppressed, material shuttled was unaffected by escitalopram and sertraline at the dose previously shown to reduce shuttling in the EBF task. Adding the extrinsic option in the DEC task appears to reduce shuttling behaviour, with lower % shuttling levels observed under these task conditions. A higher dose of sertraline was necessary to supress both total foraging and shuttling levels. As such, under these conditions, this measure may be less sensitive.

Overall, acute serotonin reuptake inhibition specifically impaired intrinsic motivation, while having limited or a positive effect on extrinsic motivation. This highlights a divergence in the pharmacological sensitivity between the two behaviours independent of task environment. Use of the individual tasks showed a broad and consistent divergence across the three compounds, while use of the DEC task has revealed further differences in behavioural effect between compounds. This may be driven by a more complex choice environment, where a wider neurobiological network is recruited to select between behavioural options. Acute increase in serotonin may exert its effect on motivation through modulation of dopamine transmission^33^. However, the direction of this relationship is heterogenous. Previous work has shown that Dorsal Raphe Nucleus stimulation increases dopamine release in the Nucleus Accumbens but supresses it in the Striatum^34^. The complexity of this relationship reveals how behaviours can be differently modulated by the same drug.

Direct pharmacological inhibition of dopamine has been utilised to establish its role in effort-related motivation. Use of the postsynaptic D_1_/D_2_ receptor antagonist haloperidol has previously been shown to decrease motivation across a range of EfR-related tasks^8,24^, as well as the newer EBF task^14^. Here, we found this effect robustly replicated in the DEC task environment, suggesting that inhibition of post-synaptic dopamine signalling reduces both types of motivated behaviour in the DEC task environment. Tetrabenazine similarly reduces monoaminergic signalling including both dopamine and serotonin via presynaptic inhibition of VMAT2. Like haloperidol, it is impairing across EfR tasks^11,35,36^ and in the EBF task. However, unlike haloperidol, it had limited effect on behavioural output in the DEC task. This suggests that motivated behaviours in this task environment are sensitive to a mechanism-specific reduction in dopamine signalling rather than an overall decline in extracellular monoamine levels. Simultaneous reduction of both serotonin and dopamine signalling may mitigate the impairing effects of a dopamine-specific reduction in signalling.

Drugs which increase dopamine signalling either through effects on the transporter or via agonism of specific receptors are of particular interest due to their potential clinical utility. The D_2_/D_3_ agonist Pramipexole, and indirect dopamine agonists and psychostimulants amphetamine and methylphenidate induced different effects in the DEC task. Here, methylphenidate increased shuttling while amphetamine impaired foraging performance in only highly motivated mice determined by a median split. This difference was previously attributed to the stimulant property of amphetamine which makes it difficult to switch between different behavioural strategies within-task, required for successful foraging^14,37^. Though a D_2_/D_3_ receptor agonist, pramipexole preferentially activates presynaptic D_2_ autoreceptors at low doses, including 0.1mg/kg used here^38^, leading to a reduction in dopamine signalling. Pramipexole reduced foraging consistent with a potential negative feedback-related decline. Contrasting previous EfR work, these compounds did not increase number of trials as expected^24^. Taken with the antidepressant findings, this measure appears less sensitive to compounds in the DEC task environment. Recent work has similarly found that altering parameters of a standard EfR task changed sensitivity and direction of effect across a range of dopaminergic compounds. Most notably, amphetamine produced opposing effects depending on task structure^39^. This highlights how certain drug effects on effortful extrinsic reward responding may only be relevant in specific contexts, and use of translatable, ethological contexts should therefore be employed wherever possible.

## Conclusion

Overall, a reduction in dopaminergic signalling had an equivalent effect across the behavioural options within the DEC task, though this effect was dependent on drug mechanism. In contrast, acute serotonergic and dopaminergic signalling increase produced a more heterogenous pattern, indicating nuanced mechanistic differences between the two types of motivation even when in the same task environment. The DEC task therefore provides a complementary approach for characterising drug effects on distinct forms of motivated behaviour within a shared environment, revealing differences that may be obscured by use of individual motivation assays. However, while it provides initial mechanistic insight, systemic pharmacology cannot distinguish direct effects on the neural processes supporting each behaviour. Given the clear complexity in our pattern of findings, future work should directly explore the mechanistic differences between the two motivated behaviours through specific neural manipulation.

## Acknowledgements

The authors would like to gratefully acknowledge Julia Bartlett for their technical assistance in behavioural testing, and the University of Bristol Animal Services Unit for care of the mice.

## Author contributions

MGJ-Conception and design of work, acquisition of data, analysis and interpretation of data, writing and editing manuscript, final approval of draft. FF-acquisition of data, analysis and interpretation of data, final approval of draft. FX— acquisition of data, analysis and interpretation of data, writing and editing manuscript, final approval of draft. CM-acquisition of data, final approval of draft. PS-acquisition of data, final approval of draft. EG-acquisition of data, final approval of draft. ESJR-writing and editing manuscript, final approval of draft.

