## Supplementary document for "A Dual Effort Choice paradigm reveals pharmacological divergence between intrinsic and extrinsic motivation within the same task environment"

| Cohort | Sex | Task manipulation | Drug | Age (weeks) | Housing | Previous exper. |
| --- | --- | --- | --- | --- | --- | --- |
| MJ20  (n=6) | M | Pre-feeding |  | 15 | Techniplast 1284 | Naive |
|  |  | Food restriction |  | 16 |  |  |
|  |  | Effort modulation (EBF) |  | 18 |  |  |
|  |  | Effort modulation (FED) |  | 20 |  |  |
| ME1+2  (n=16) | M+F |  | Escitalopram | 20 | Techniplast 1284 | EBF-rivastigmine |
|  |  |  | Methylphenidate | 24 |  |  |
| MJ27+  28  (n=16) | M+F |  | Sertraline (3mg) | 14 | Techniplast 1284 | Naive |
|  |  |  | Venlafaxine | 16 |  |  |
|  |  |  | Sertraline (10mg) | 18 |  |  |
|  |  |  | Tetrabenazine | 26 |  |  |
|  |  |  | Haloperidol | 27 |  |  |
| MJ30+31  (n=16) | M+F |  | Amphetamine | 12 | Techniplast **XXXX** | Naive |
|  |  |  | Pramipexole | 14 |  |  |

**S2**

Enrichment in the 1284 cages consisted of a paper shelter, wooden chew, aspen ball, small cardboard tube, large cardboard tube, wooden labyrinth and ceiling tube. Mice had paper nesting material and woodchip bedding. The 1290 cages consisted of the same enrichment with the addition of a Nombrero (Datesand, UK) hanging from the ceiling (a rat treat holder repurposed as a platform for mice to swing from) and two T-bricks (Datesand, UK) fixed to the ceiling, allowing for climbing across platforms.

**S3**

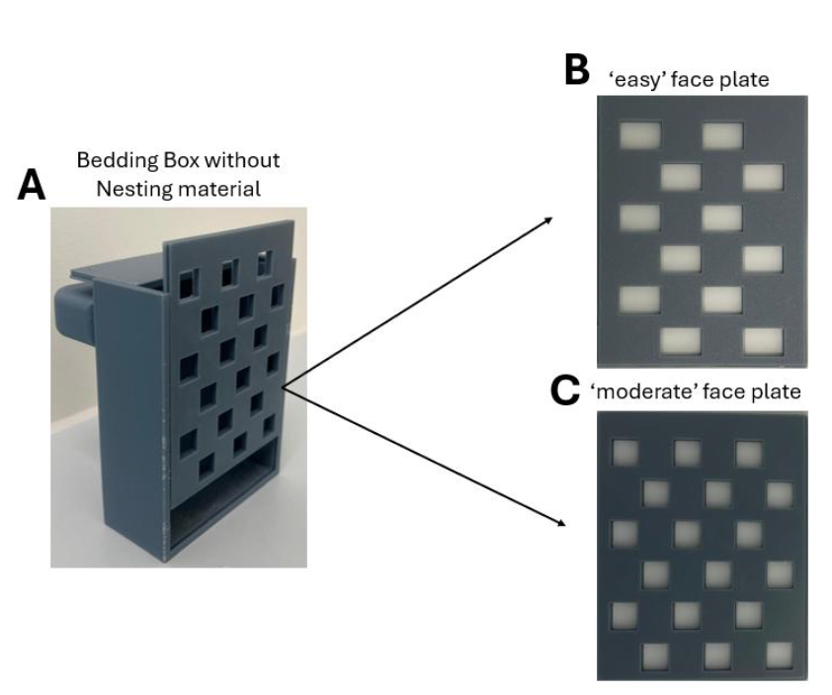

***S3. Nesting material box. A*** *nesting material box comprises a main body, face plate, lid and magnet bar.* ***B*** *face plate with 1.5cm^2^ apertures represent the ‘easy’ difficulty and* ***C*** *the 0.75cm^2^ apertures represent the ‘difficult’ difficulty.*

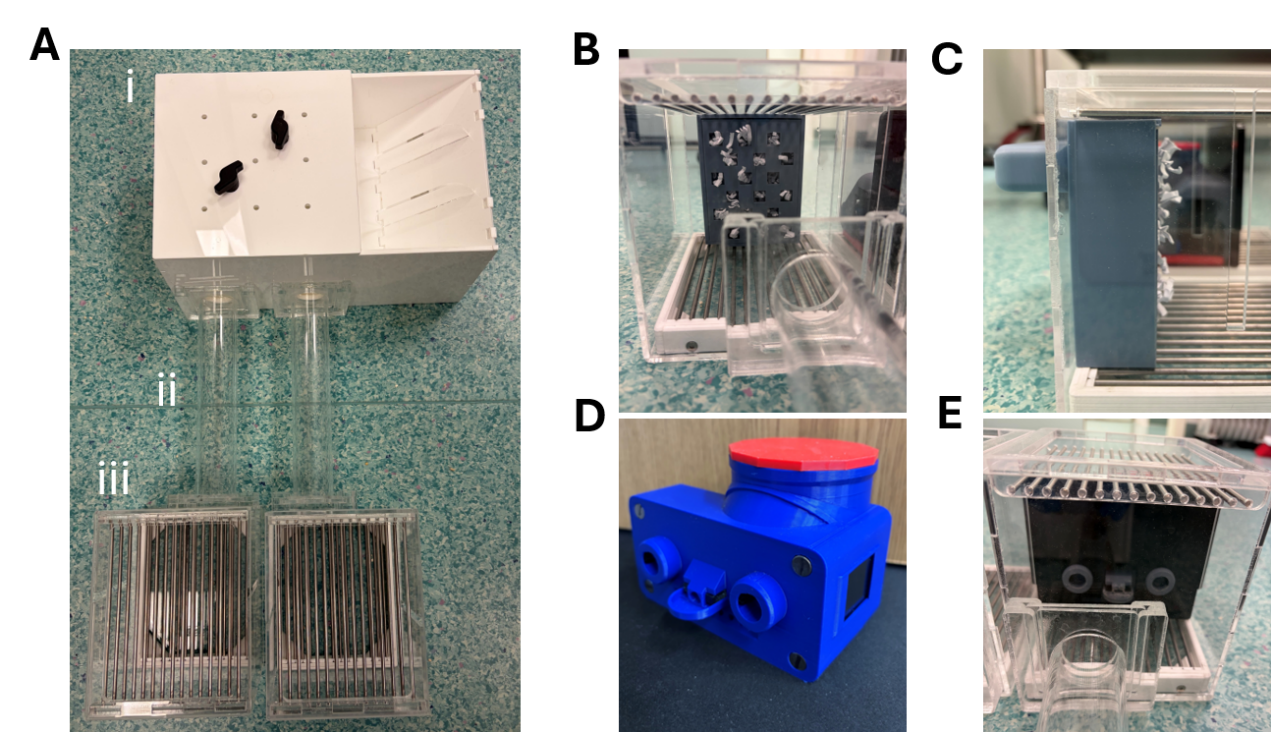
**S4**

***S4. Dual Effort Choice task apparatus. A*** *Full task apparatus for the Dual Effort Choice task. The arena comprises (i) an enclosed home-like area with woodchip at the base, a bowl for food and water bottle mount. The arena is covered by a removable lid with small holes to allow some air circulation. The mice can freely choose to shuttle through clear tubes (ii) to access one of two separate compartments (iii) that separately contain the FED device and the nesting material box. The compartments contain a barred floor and a barred ceiling to promote air flow and create a more ‘open’ environment.* ***B*** *Sizzle nesting material is accessed through apertures in a 3D printed nesting material box fixed to the back of the compartment. Mice can freely choose to pull nesting material from the box using their teeth. Mice can then choose to shuttle the nesting material through the tube to the home area, or drop it out in the open. The apertures in the picture depict the moderate (1cm^2^) apertures used throughout the pharmacology experiments.* ***C*** *The nesting material box is fixed to the back of the arena using an external magnet bar which prevents it from falling when the mouse pulls the material.* ***D*** *The FED device comprises of two nose poke apertures either side of a reward delivery magazine. The FED dispenses a 20mg reward pellet upon a set number of nose pokes into the correct aperture.* ***E*** *The FED device is protected by a Perspex screen which allows access to the nose pokes and reward magazine only.*

**S5**

Before onset of experimentation, mice were habituated to a 20% condensed milk solution (Nestlé Carnation mixed with tap water) in the home cage as described on the 3Hs Initiative website. Next, mice were given 300µl (if male) or 200µl (if female) via a 1ml syringe placed through the bars of the home cage. The syringe was depressed slowly, matching the drinking rhythm of the mouse. This was repeated once per day until the mouse rapidly approached consumed the syringe contents (~approximately three sessions).

**S6**

Mice were first habituated to a piece of Vetbed, a soft piece of material on which dosing will take place. In accordance with 3Hs protocols, mice are first allowed to freely explore for 10 seconds before returning to the home cage. A palatable reward (20mg reward pellets) is put in the home cage immediately after Vetbed exposure to promote a positive association with the Vetbed. On the second session, the mouse is covered loosely with a hand for a few seconds. On the third day, the mouse is held more firmly, covering the mouse with the palm (head pointing towards the wrist). On the final day, the mouse is covered firmly again, and this time flank fat is held between two fingers, creating a tent for injection. Upon the release, the mouse should not dart away and should be easily cupped back into the cage. At this point, the mouse is habituated and suitable for dosing.

**S7**

| **Experiment** | **Measure** | **Statistical exclusion in dataset** |
| --- | --- | --- |
| **Escitalopram**  ***EBF option***  ***FED3*** | Total foraged | N=1 vehicle, n=1 3mg/kg outliers |
|  | Total taken through | N=1 vehicle, n=1 3mg/kg outliers |
|  | %taken through | N=1 vehicle outlier |
|  | Number of trials | N=2 pre-excluded due to jammed devices, n=1 vehicle outlier |
| **Venlafaxine**  ***EBF option***  FED3 | Total foraged | N=1 excluded outlier in 3 out of 4 measures, n=1 10mg/kg outlier |
|  | Total taken through | N=1 vehicle, n=1 10mg/kg outliers |
|  | %taken through | No exclusions |
|  | Number of trials | N=1 vehicle outlier |
| **Sertraline 3mg/kg**  ***EBF option***    ***FED3*** | Total foraged | N=2 3mg/kg outliers |
|  | Total taken through | N=1 vehicle, n=1 3mg/kg outliers |
|  | %taken through | No exclusions |
|  | Number of trials | N=1 vehicle, n=1 3mg/kg outliers |
| **Sertraline 10mg/kg**  ***EBF option***  ***FED3*** | Total foraged | N=1 vehicle outlier |
|  | Total taken through | N=1 excluded due to 2 outlying values |
|  | %taken through | N=1 vehicle outlier |
|  | Number of trials | N=1 10mg/kg outlier |
| **Amphetamine**  ***EBF option***  ***FED3*** | Total foraged | N=1 vehicle, n=1 0.3mg/kg outlier |
|  | Total taken through | N=1 excluded due to 2 outlying values |
|  | %taken through | No exclusion |
|  | Number of trials | N=1 0.3mg/kg outlier |
| **Methylphenidate**  ***EBF option***  ***FED3*** | Total foraged | N=1 1mg/kg, n=2 3mg/kg outliers |
|  | Total taken through | N=2 vehicle, n=1 1mg/k, n=1 3mg/kg, n=1 10mg/kg outliers |
|  | %taken through | N=1 vehicle, n=1 1mg/kg. n=1 10mg/kg outliers |
|  | Number of trials | N=2 pre-excluded due to jammed devices, n=1 3mg/kg, n=1 10mg/kg outliers |
| **Tetrabenazine**  ***EBF option***  ***FED*** | Total foraged | N=1 excluded due to 2 outlying values |
|  | Total taken through | N=1 vehicle, n=1 6mg/kg, outliers |
|  | %taken through | No exclusion |
|  | Number of trials | N=1 vehicle outlier |
| **Haloperidol**  ***EBF option***  ***FED3*** | Total foraged | N=1 scheduled 1, n=1 vehicle outlier |
|  | Total taken through | N=1 vehicle outlier |
|  | %taken through | Not statistically analysed due to lack of foraging. |
|  | Number of trials | N=1 vehicle outlier |
| **Pramipexole**  ***EBF option***  ***FED3*** | Total foraged | N=1 0.1mg/kg outlier |
|  | Total taken through | N=1 0.1mg/kg outlier |
|  | %taken through | No exclusion |
|  | Number of trials | N=1 vehicle, n=1 0.1mg/kg outliers |
| ***Pramipexole***  ***EBF task*** | Total foraged | N=2 vehicle, n=2 0.001mg/kg, n=2 0.003mg/kg n=2 0.1mg/kg outliers |
|  | Total taken through | N=2 vehicle, n=2 0.001mg/kg, n=2 0.003mg/kg n=2 0.1mg/kg outliers |
|  | %taken through | N=2 vehicle, n=2 0.001mg/kg, n=2 0.003mg/kg n=2 0.1mg/kg outliers |
| ***Tetrabenazine***  ***EBF task*** | Total foraged | N=1 excluded due to 3 out of 5 outlying values, n=1 6mg/kg outlier |
|  | Total taken through | N=1 excluded due to 3 out of 5 outlying values, n=1 6mg/kg outlier |
|  | %taken through | N=1 2mg/kg, n=1 4mg/kg, n=1 6mg/kg, n=1 8mg/kg outliers |
|  | General locomotor activity | N=2 vehicle, n=4 2mg/kg, n=2 4mg/kg, n=4 6mg/kg, n=4 8mg/kg missing values |

***Table of outliers and exclusions.***

**S8**

There was a main effect of pramipexole on total nesting material foraged (F _(2.313, 34.70)_ = 3.290, p = 0.0427). Post-hoc comparisons didn’t reach significance (**S8A**). There was a main effect of pramipexole on total nesting material shuttled (F _(2.370, 35.56)_ = 4.556, p = 0 .0131). 0.1mg/kg pramipexole caused a reduction in nesting material shuttled compared to the vehicle (p=0.0439). There was a trend towards decrease in mice treated with 0.03mg/kg too (p=0.0914) (**S8B**). There was a main effect of the drug on % nesting material shuttled (F _(1.984, 29.76)_ = 3.389, p = 0.0476). Post-hoc comparisons didn’t reach significance (**S8C**).

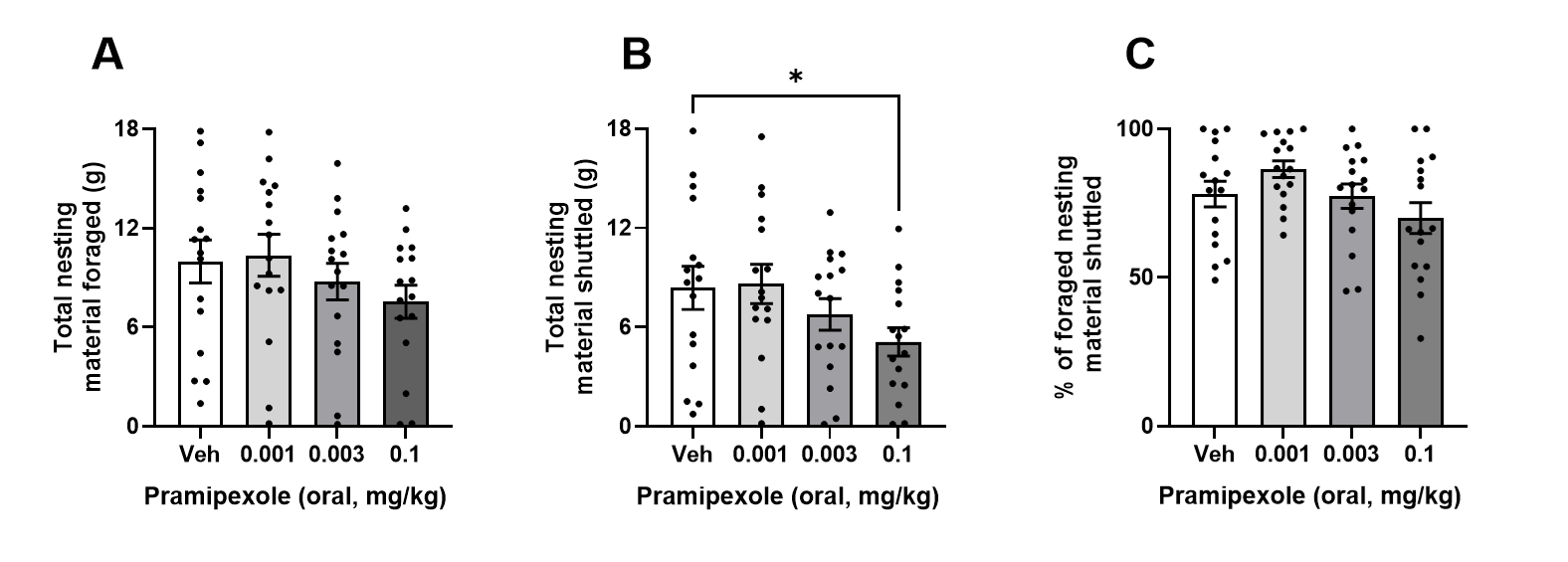

***S8. Pramipexole reduced shuttling in the EBF task. A*** *Pramipexole did not affect foraging but did impair shuttling* ***B*** *at 0.1mg/kg.* ***C*** *% shuttled was unaffected. *p<0.05, bars are mean ± SEM with data points overlaid.*

**S9**

There was a main effect of tetrabenazine on total nesting material foraged (F_(3.304, 46.26)_ = 9.592, p < 0.0001), where mice treated with 6 and 8mg/kg tetrabenazine foraged less compared to vehicle (p=0.0168, p=0.0021 respectively) (**S9A**). There was a main effect of tetrabenazine on total nesting material shuttled back to the main box (F _(3.467, 48.54)_ = 6.843, p = 0 .0003), where mice treated with 6 and 8mg/kg tetrabenazine shuttled less compared to the vehicle (p=0.0409, p=0.0019 respectively) (**S9B**). There was no effect of the drug on % nesting material shuttled (**S9C**). Tetrabenazine had a main effect on forage area movement (F _2.424, 26.66)_ = 4.813, p = 0.0121). Movement was measured by a passive infrared sensor system positioned over the forage area of the EBF arena. 8mg/kg tetrabenazine reduced movement in the forage area (p=0.0065) (**S9D**). Since 8mg/g of the drug had an effect not only on motivation to perform the task but on movement as well 6mg/kg was chosen to be tested on the DECT task.

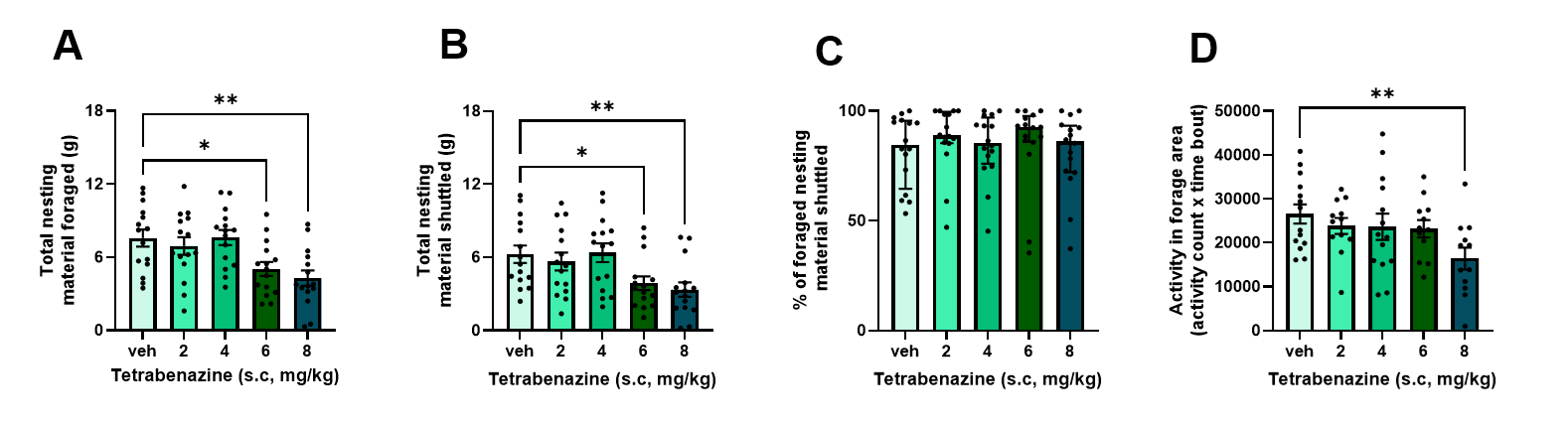

***S9. Tetrabenazine reduced foraging and shuttling in the EBF task. A*** *Tetrabenazine reduced foraging at 6 and 8mg/kg* ***B*** *as well as shuttling.* ***C*** *% shuttled was unaffected.* ***D*** *Tetrabenazine reduced movement in the foraging area at 8mg/kg. *p<0.05, **P<0.01, bars are mean ± SEM with data points overlaid.*
